# Replication of single-cell proteomics data reveals important computational challenges

**DOI:** 10.1101/2021.04.12.439408

**Authors:** Christophe Vanderaa, Laurent Gatto

## Abstract

**Introduction:** Mass spectrometry-based proteomics is actively embracing quantitative, single-cell level analyses. Indeed, recent advances in sample preparation and mass spectrometry (MS) have enabled the emergence of quantitative MS-based single-cell proteomics (SCP). While exciting and promising, SCP still has many rough edges. The current analysis workflows are custom and built from scratch. The field is therefore craving for standardized software that promotes principled and reproducible SCP data analyses.

**Areas covered:** This special report is the first step toward the formalization and standardization of SCP data analysis. scp, the software that accompanies this work, successfully replicates one of the landmark SCP studies and is applicable to other experiments and designs. We created a repository containing the replicated workflow with comprehensive documentation in order to favor further dissemination and improvements of SCP data analyses.

**Expert opinion:** Replicating SCP data analyses uncovers important challenges in SCP data analysis. We describe two such challenges in detail: batch correction and data missingness. We provide the current state-of-the-art and illustrate the associated limitations. We also highlight the intimate dependence that exists between batch effects and data missingness and offer avenues for dealing with these exciting challenges.

**Article highlights:**

- Single-cell proteomics (SCP) is emerging thanks to several recent technological advances, but further progress is still lagging due to the lack of principled and systematic data analysis.
- This work offers a standardized solution for the processing of SCP data demonstrated by the replication of a landmark SCP work.
- Two important challenges remain: batch effects and data missingness. Furthermore, these challenges are not independent and therefore need to be modeled simultaneously.

## 2 Introduction

High-throughput single-cell assays are instrumental in highlighting the biology of heterogeneous cell populations, tissues and cell differentiation processes. Single cell RNA sequencing (scRNA-seq) is a prominent player, thanks to its throughput, technical diversity, and the computational tools that support its analysis and interpretation. scRNA-seq is however blind to the many biologically active gene products, proteins and their many proteoforms. Mass spectrometry (MS)-based approaches to study the proteome of single cells are emerging, using the wide range of possibilities offered by the technology, including miniaturized and automated sample preparation, labeled and label-free quantitation, as well as data dependent and independent approaches [1, 2, 3, 4, 5]. All these avenues promise to be valuable contributions to the single cell tool kit.

In this work, we focus on the processing of mass spectrometry-based single cell quantitative data, as produced from the raw data using widely used tools such as, for example, MaxQuant [6] or Proteome Discoverer (Thermo Fisher Scientific). As expected for a young and fast evolving field such as single-cell proteomics (SCP), there are yet no best practice nor any consensus as to how to adequately process such data. Some studies start from protein and peptide tables as produced by MaxQuant followed by manual data manipulation using Excel [7, 8], proceed with Perseus [9, 10], use private in-house scripts [11, 7, 12], while others publish their custom scripts openly [13, 14]. In this report, we present the replication of the open-source scripts of SCoPE2 published by Specht and colleagues and their implementation as a formal R/Bioconductor package named scp [14, 15]. We have chosen to focus on the replication of the SCoPE2 analysis for several reasons. First, it put a milestone in the SCP field by reporting the acquisition of over a thousand single cells and proving that SCP has reached its potential in becoming a high-throughput technology [16].

Furthermore, the acquisitions were distributed across several MS runs, chromatographic batches and 2 labeling protocols, offering the opportunity to assess confounding effects that are common in complex experimental or clinical designs. Finally, the authors publicly shared all the code and data necessary to reproduce the results presented in their article, making benchmarking of the replication possible. Replicating their work allows to formalize and standardize the current SCP data processing pipeline, and also brings to light two important challenges for SCP data analysis that we will address in the Expert Opinion section.

## 3 Replicating the SCoPE2 analysis

We focused on replicating the SCoPE2 analysis by Specht et al. [14] since their raw and quantitative data and the processing scripts are readily available, making this work an outstanding example of open science. Sharing research outputs is indeed paramount to promote community-wide contributions that will further push the development of the field forward. Furthermore, Specht et al. implemented new metrics and quality controls that the field could benefit from. Although the provided data and scripts could fully reproduce their results, the code is difficult to read for non expert programmers and lacks modularity, making it tedious to reuse and hard to adapt and extend. We therefore decided to provide a standardized and modularized framework to replicate this analysis and hence offer a common ground for SCP data analysis and method development. Our data structure relies on two established R/Bioconductor [17] data classes: `QFeatures` [18] and `SingleCellExperiment` [19]. `QFeatures` is a data object model dedicated to the manipulation and processing of MS-based quantitative data. It explicitly records the successive steps to allow users to navigate up and down the different MS levels. `SingleCellExperiment` is another efficient data container, that serves as an interface to state-of-the-art methods and algorithms for single-cell data. Our approach combines these two classes and benefit from their respective strengths. Based on this data framework, we built two pieces of software: `scp` and `scpdata`.

The `scp` package extends the functionality of `QFeatures` to SCP applications (Figure 1A, orange boxes). It includes functionality that was implemented in SCoPE2, such as normalization by a reference channel, filtering single-cells based on the median coefficient of variation, or filtering of peptide-spectrum matches (PSM) based on the single-cell to carrier ratio (SCR). A core feature of the `scp` package is the conversion of standard data tables, like those exported by MaxQuant or Proteome Discover (Thermo Fisher Scientific), to `scp` formatted data objects along with sample metadata. `scpdata` disseminates twelve SCP data sets formatted using our data structure. The purpose of `scpdata` is three-fold. First, it is an ideal platform for data sharing and hence lays the ground for open and reproducible science in SCP. For instance, the package provides, among others, the PSM, peptide and protein data used for this replication study. Second, it facilitates the access to SCP data for developers to build and benchmark new methodologies. Finally, the `scpdata` package allows new users to readily access curated and thoroughly annotated SCP data in the context of training and education.

**Figure 1:**
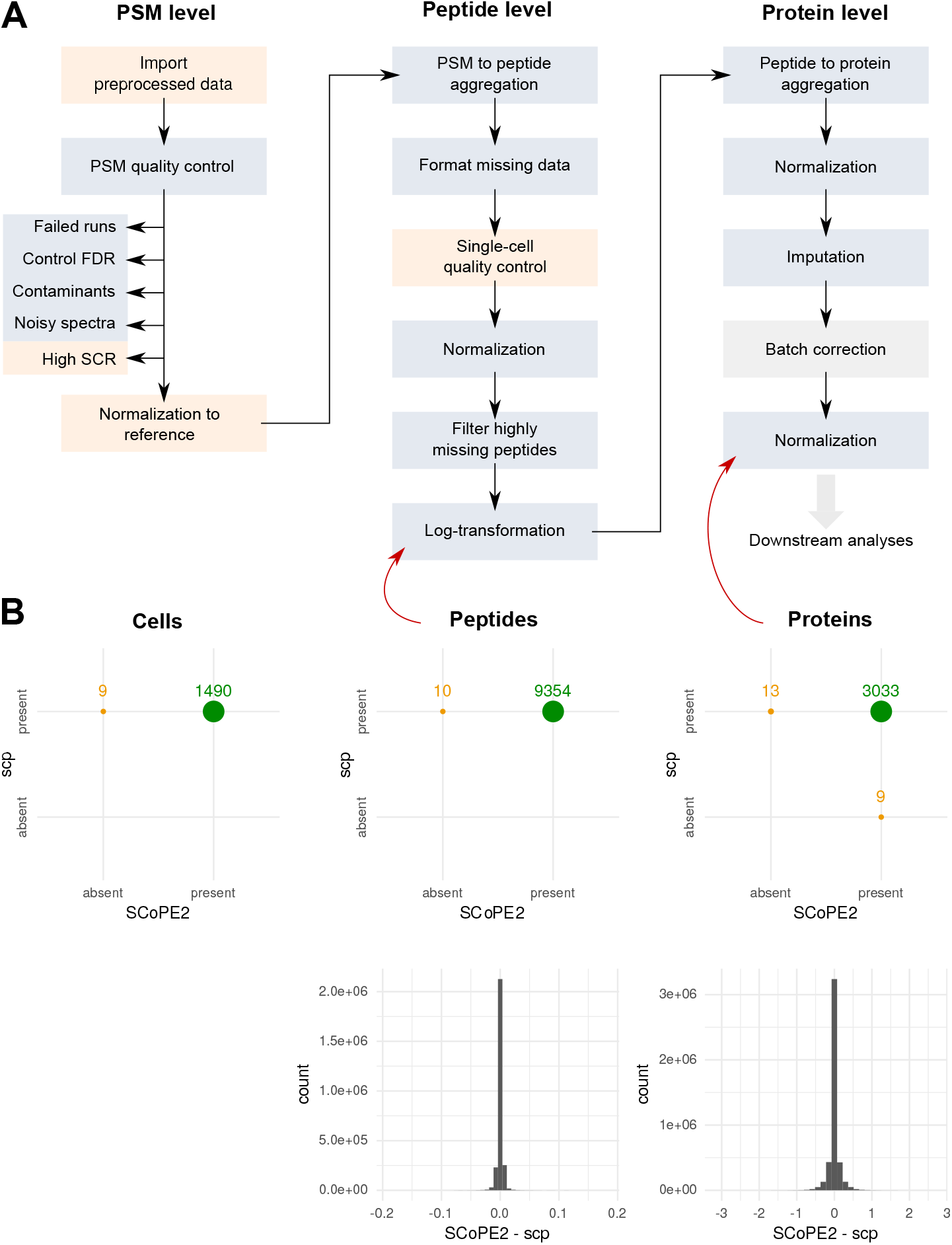
Replication the SCoPE2 data processing. **A:** Overview of the key steps performed in the SCoPE2 script. Blue boxes indicate steps that were already available in `QFeatures`. The orange boxes indicate steps that were implemented in `scp`. The gray box indicates a step implemented in another package. **B:** Results of the replication. The top row demonstrates the agreement between the number of cells, peptides or proteins obtained when running the original SCoPE2 scripts and `scp`. The bottom row shows the numerical differences between the peptide or protein expression matrices. Red arrows point towards the step that generated the tested data.

The first step of the replication was to retrieve the SCoPE2 data. The data are hosted on Google Drive and are clearly linked from the authors’ web page [20]. We formatted the data set using `scp` and included it in `scpdata` along with comprehensive documentation about data content, data acquisition and data collection. This is true for any data set in `scpdata`. Next, we retrieved the SCoPE2 code from the authors’ GitHub repository and formalized the key steps of the workflow (Figure 1A) [20]. Most steps implemented in SCoPE2 are routinely performed in bulk proteomics and are handled by existing software such as `QFeatures`. The reuse of existing code is essential in software development because it allows the developer to focus on the innovative aspects of its research field without losing time reinventing the wheel [17]. Next, we implemented the missing steps in `scp` (Figure 1A, orange boxes) and provided clear documentation and examples. Finally, we wrote a new workflow that fully replicates the results of the SCoPE2 script, using our standardized software. The output obtained after running the `scp` workflow leads to very similar results compared to the data provided by the authors (Figure 1B). The sets of filtered cells and proteins are almost identical. The final processed data using the two workflows show high similarity with most differences close to zero. A small proportion of the processed protein expression values show larger differences between the two workflows. These values arise during imputation of the protein data that will be discussed in a later section (Figure 4B). A detailed report about the replication of the SCoPE2 analysis using scp can be found on GitHub [21]. This report includes the code used and some comprehensive documentation to give readers a good understanding of the underlying processes. It demonstrates how to integrate other tools to the workflow, such as the `ggplot2` package for data visualization [22]. It also gives additional comments on each steps of the SCoPE2 workflow and suggests alternatives steps and methods for future analyses.

## 4 Conclusion

New tools are required for principled and standardized analysis of SCP data. In this work, we show the successful application of scp, our R/Bioconductor software package, to replicate the data processing workflow published in [14]. While replication or reproduction don’t guarantee optimal processing of the data and the validity of the results, they demonstrate coherence and increase trust in the data and the results. In addition, the `scp` package allows for an open SCP environment that can foster new methodological developments as well as spreading SCP data analysis towards a broader computational community. We emphasized the standardization of the implementation which facilitates the integration with currently available tools such as the single-cell methods and workflows provided by the Bioconductor project [19]. Furthermore, the code is continuously tested and improved to guarantee long term usability of the software.

Although the replication of the SCoPE2 results supports the reliability of the original work, additional improvements are necessary. Complex challenges, such as batch effects and data missingness still need to be tackled and further methodological developments are required to obtain an optimal workflow.

## 5 Expert opinion

### 5.1 Batch correction

The SCoPE2 protocol relies on sample multiplexing. The 1490 single-cell samples were multiplexed across 177 MS acquisitions, 63 of which were labeled using TMT-11 and 14 using TMT-16. The data were acquired across 4 chromatographic batches (LCA9, LCA10, LCB3 and LCB7). Unsurprisingly, batch effects account for the main source of variation in the unprocessed peptide data, as indicated by a principal component analysis (PCA) on Figure 2. The first component (12.4 % of total variance) perfectly separates the TMT-11 from the TMT-16 batches and the second component (6 % of total variance) further separates the four chromatographic batches. The next two components (7.3 % of total variance) are driven by biological variations and separate macrophages from monocytes. Because components in PCA are constrained to be orthogonal, this analysis indicates that most of the technical and biological variations are independent. This is a key assumption in order to separate the undesired technical variability from the biological variability. Orthogonality between technical and biological variation is achieved by carefully designing the experiment. As pointed out in the SCoPE2 protocol, it is crucial to randomize cell types and biological samples across different MS batches.

**Figure 2:**
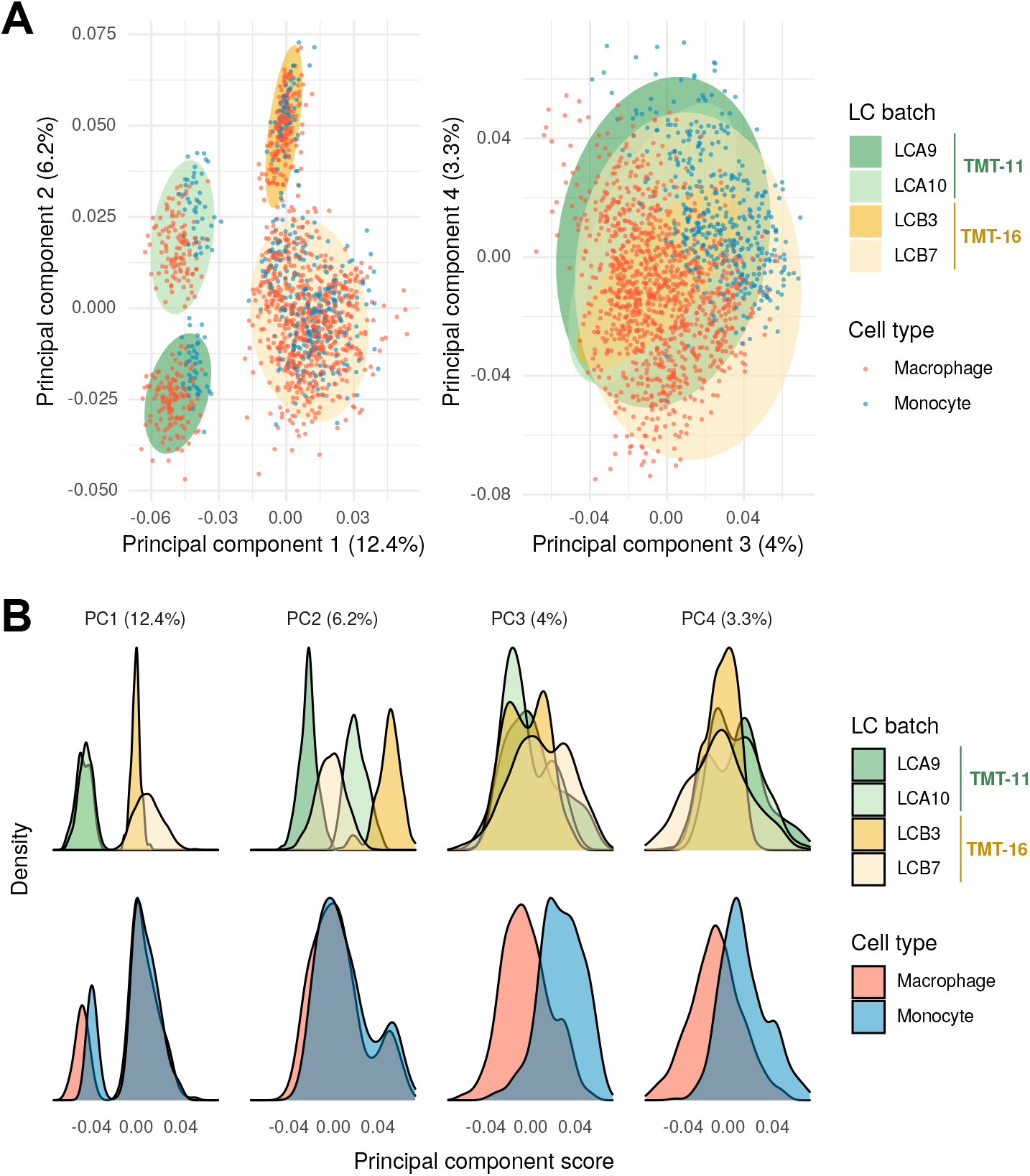
SCP data exhibit batch effects. The PCA is performed on the peptide data after log-transformation (*cf* Figure 1A). The Nonlinear Iterative Partial Least Squares (NIPALS) algorithm was used to account for missing values during PCA [23]. The liquid chromatography (LC) batches were acquired either with a TMT-11 (green ellipses) or TMT-16 (yellow ellipses) protocol. The data set contains two types of single-cells: macrophages (red dots) and monocytes (blue dots). **A:** PCA scores for the first four components. Each point represents a single cell and is colored according to the corresponding cell type. The ellipses give the 95 % interval for each chromatographic batch. **B:** Distribution of the principal components scores. Each principal component is displayed in a separate column. The distributions are split according to LC batch (top row) or to the sample type (bottom row). The densities were computed from the PCA scores.

Since batch effects are technically unavoidable, they need to be accounted for computationally. The SCoPE2 authors opted for removing the batch effect using ComBat, an empirical Bayes framework [24]. As with any procedure, it is important to understand and apply the requirements of the method. First, ComBat assumes a balanced design, i.e. it requires that differences between batches be only the result of technical differences. This can be an issue when cell types or cell states are unknown in advance, for example in experiment designed to discover new cell populations. Second, ComBat cannot work with missing data, which requires the data to be imputed beforehand. As we will discuss later, imputation is a sensitive step that can lead to substantial artifacts in the data, especially when the number of missing values is high, as is the case for single-cell proteomics data. Thirdly, ComBat cannot account for the hierarchical structure of batch effects. We anticipate that once the technology matures, and is applied to clinical samples, for instance across multiple patients and acquisitions, such a hierarchical structure will become significant. Finally, ComBat creates a new data set by fitting and removing the batch effect from the input data and ignores the uncertainty associated to the estimation of the batch effect itself. It would be important to quantify this uncertainty instead of only considering point estimates. Other batch correction methods have been developed for scRNA-Seq data and were extensively benchmarked elsewhere [25, 26]. However, methods tailored for other single-cell applications only partly address the above listed issues and none suggest to propagate the uncertainty linked to batch effect estimation. An alternative approach would be to avoid batch correction altogether and account for batch effects explicitly during data modeling [27, 28, 29].

### 5.2 Data missingness

Next to batch effects, missing values are a major challenge in MS-based proteomics [30, 31, 32]. Missingness refers to the fact that not all features (peptides or proteins) are detected and quantified in all samples. We can distinguish between to types of missingness.

The first type is biological missingness. The peptides of a protein are not detected in a sample because that sample does not express the protein. Such missingness is biologically relevant and must be considered accordingly. We observed this phenomenon in the SCoPE2 data, where some peptides are systematically missing less in macrophages compared to monocytes and the reduced missingness is correlated with an increased average expression level in that cell type (Figure 3A).

**Figure 3:**
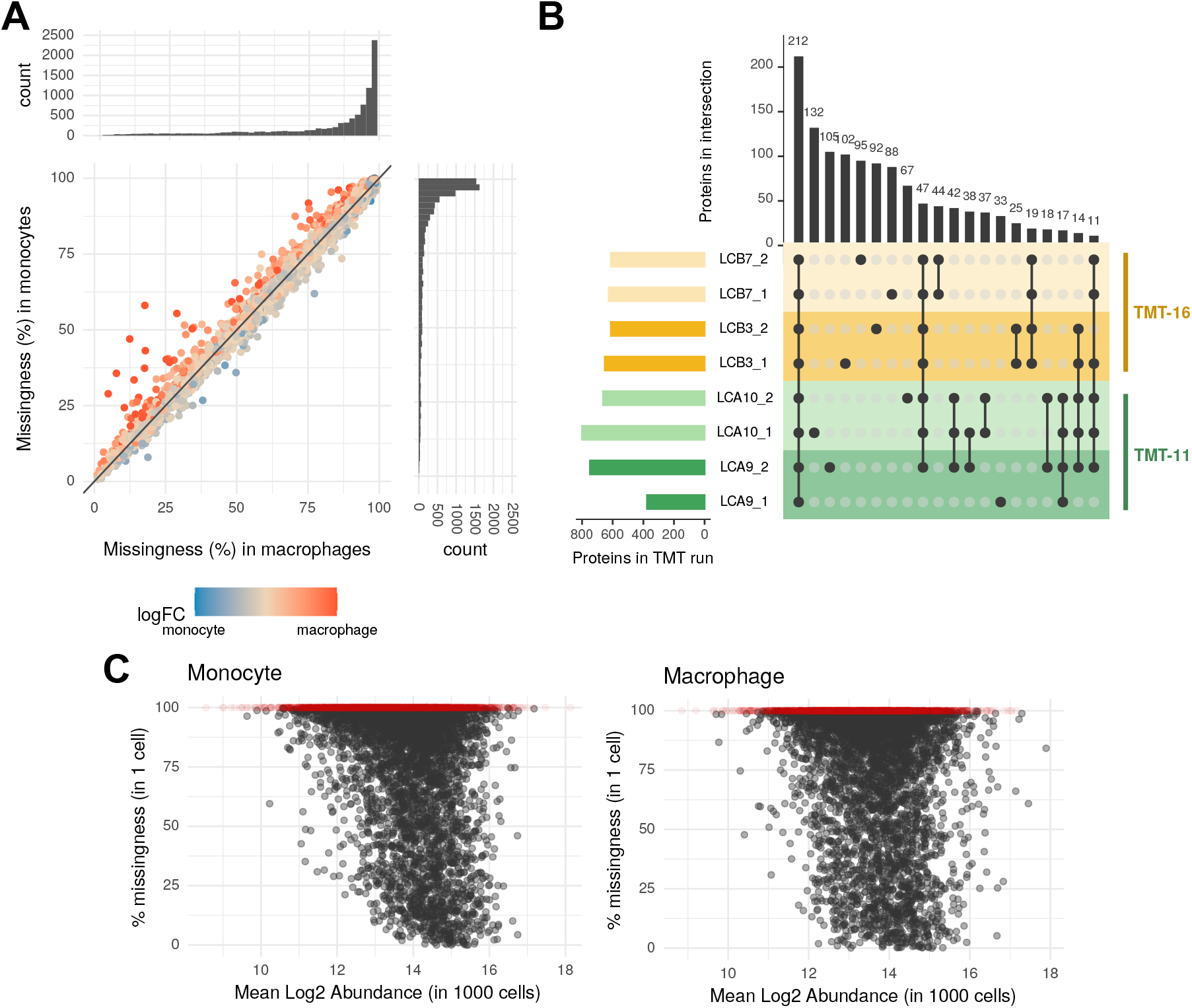
Missing data is the consequence of two components. **A.** Biological missingness is illustrated by plotting the proportion of missing values in monocytes against the proportion of missing values in macrophages for each peptide. Those proportions are also shown on the histograms along the y and x axis for monocytes and macrophages, respectively. Each peptide is colored according to its relative log fold change between macrophages over monocytes. The data used are the peptides after log-transformation (cf Figure 1A). **B.** Technical missingness between replicates is shown using an upset plot on eight representative MS runs [33]. Two MS runs were randomly sampled from each of the four LC batches. The bar plot on the left shows the total number of proteins per MS run and the bar plot at the top shows the number of proteins for each intersection. Black dots indicate which MS runs are included in the intersection. **C.** Relationship between the proportion of missingness in single-cell samples and the average log2 abundances. Each dot represents a peptide. Samples containing 1000 cells were used as a proxy to estimate the peptide abundances for monocyte (left) and macrophage (right) populations. Red dots highlight peptides that are found in 1000-cell samples but absent in single-cell samples.

The second type is technical missingness. There are several technical mechanisms that explain why a protein could not be detected in a sample. A first reason is that none of its constituting peptides could be correctly delivered to the MS instrument, for example due to sample loss. Sample loss is a major concern for single-cell applications because only limited amounts of material are available to start with. This limitation has been actively researched and improved in the last two years, and the SCoPE2 protocol or the cellenONE’s proteoCHIP are two prime examples [14, 5]. Poor ionization of peptides can also lead to reduced signal or to missing data. Another cause of technical missingness is related to MS1 peak selection. In data dependent acquisition (DDA), only the most abundant precursor peaks are selected for fragmentation and MS2 analysis. Whether a peak will be selected is therefore dependent on the abundance of the peptide and the surrounding peptides in a specific chromatographic region, as well as their ionization efficiencies. Several approaches have been developed to reduce this bias by propagating spectrum identifications from one sample to the corresponding MS1 peaks from another sample. The match between run algorithm of MaxQuant is very popular in label-free SCP [10, 9, 7, 8, 34, 12], but methodological improvements have recently been suggested for both label-free and TMT-based SCP [35, 36]. Finally, another reason for missingness is the inability to match a spectrum to a peptide sequence due to poor spectrum quality. Low-quality spectra occur because of peptide co-isolation after MS1 selection or when the peptide abundance is close to the detection limit. This limitation is tackled by improving the current sensitivity of LC-MS/MS instruments. For instance, [8] reported an increased proteome coverage when decreasing the diameter of the LC columns for improved chromatographic resolution or upgrading the Orbitrap Eclipse Tribrid MS to an Orbitrap Fusion Lumos Tribrid MS for improved MS sensitivity. Later, they also showed improved peptide identification by coupling the MS with a high field asymmetric ion mobility spectrometry (FAIMS) device [34]. Technical missingness translates to the fact that two similar MS runs will not contain the same set of quantified peptides and proteins. Although most proteins are common to several MS runs, each run exhibits a specific set of proteins that were probably present but missed in the other runs (Figure 3B). We observe only marginal correlation between the peptide abundance and the proportion of missingness in single-cell samples (Figure 3C), as was already described for bulk proteomics [31]. While upcoming technical improvements to SCP will further decrease the amount of missing values, computational approaches will still be required.

To overcome the current limitations regarding missing data, Specht and colleagues imputed missing data using the k-nearest neighbors (KNN) method. They applied KNN in the sample space instead of the gene space, thus increasing the similarity between samples. Since subsequent cluster or differential abundant protein analyses focus on sample-wise differences, this causes an underestimation of the variance and hence leads to a potential increase of false positive outcome.

Furthermore, the imputation is performed at the protein level. As pointed out by [30], imputation at the protein level means that a first implicit imputation is performed at the peptide level and the authors instead suggest to use well-justified imputation methods directly at the peptide level. However, a good understanding of the missingness mechanism is required to justify the use of a suited imputation method. Further research is required to extend current work on bulk proteomics to the context of SCP data [30, 31]. Finally, just like batch correction, imputation is an estimation process that generates estimates with some degree of uncertainty. Replacing missing data by imputed values ignores the variance associated to the estimates and this variance can become large when available data are scarce. Multiple imputation, i.e. the application of a range of imputation parameters or methods to estimate a range of plausible values rather than point estimates, would be a promising strategy here. This is best illustrated by an issue we noticed in the data. For instance, the E3 ubiquitin-protein ligase (RNF41) is quantified in only three MS runs and KNN imputation predicted the missing values for the remaining runs (Figure 4A). When comparing the resulting data distribution to the distribution for vimentin (VIM), a protein that is not missing, we can clearly observe that the imputation introduces two suspicious trends. First, the variability observed for imputed values is much lower than for acquired values, and second, the imputation does not exhibit batch effects. While reduced variability and absence of batch effects are desirable properties, in this case, we are faced with erroneous data that does not hold biologically meaningful information. The imputed data for RNF41 is unreliable and should be flagged accordingly. Furthermore, small quantification errors may get amplified during the imputation step, as observed during the replication of the SCoPE2 data (Figure 4B). While we observed minimal differences between the two workflows before imputation, those small differences where magnified after imputation, considerably increasing the proportion of values deviating from zero.

**Figure 4:**
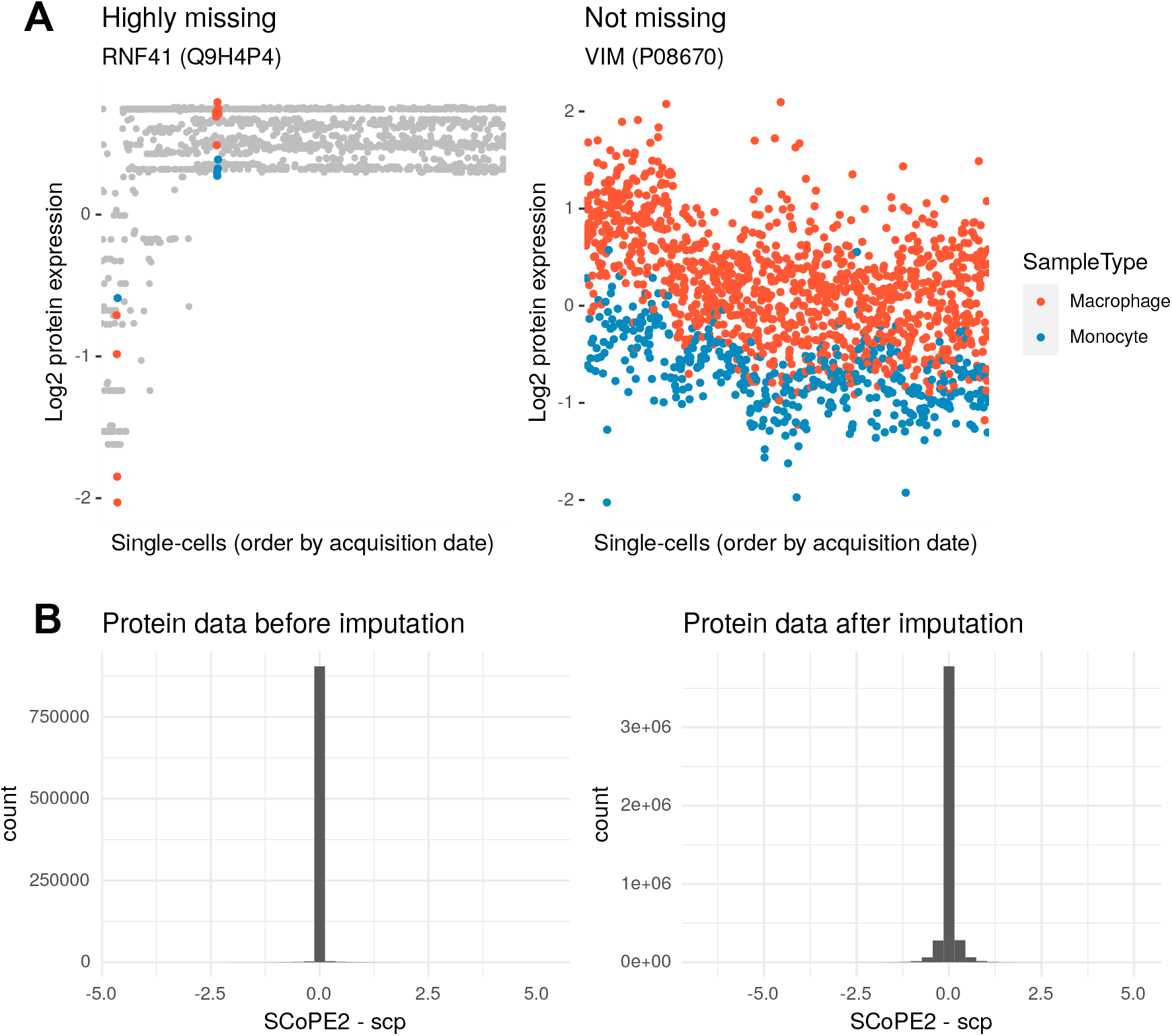
Problem with imputation. **A.** The data distribution is shown for two proteins: RNF41, a highly missing protein, and VIM, a protein with no missing data. The variance associated to the imputed values for RNF41 (gray points) is not correctly estimated as compared to the variance observed for VIM. Data points are colored in red for macrophages and in blue for monocytes. **B.** Numerical differences between the protein data generated by the scp and the SCoPE2 processing workflows, before imputation (left, *cf.* Figure 1A, *Normalization* step) and after imputation (right, *cf.* Figure 1A, *Imputation* step).

### 5.3 Batch effects and data missingness are not independent

As of today, all published SCP analyses consider batch effects and data missingness as two distinct issues that can be tackled separately when they are, in reality, correlated. Figure 5 highlights the impact of acquiring data in different LC batches on the data missingness. First, since with SCoPE2, peptides are identified from the carrier signal, the number of identified peptides and their missingness display a prominent MS acquisition effect. Second, the LC batches influence the amount of missingness. For instance, more missing values are observed for LCB3 than LCB7. Third, the amount of missing data within each LC batch varies over time. LCA10 displays a very clear increase of missingness, while LCB7 shows a decrease. Finally, LC batches also influence the variability of missing data as the proportion of missing values both within and between MS runs, with LCB3 displaying much less thereof compared to all other ones. Therefore missing values can only be correctly modeled if we include batch covariates. Inversely, batch effect can only be correctly modeled if we accurately model the missing data.

**Figure 5:**
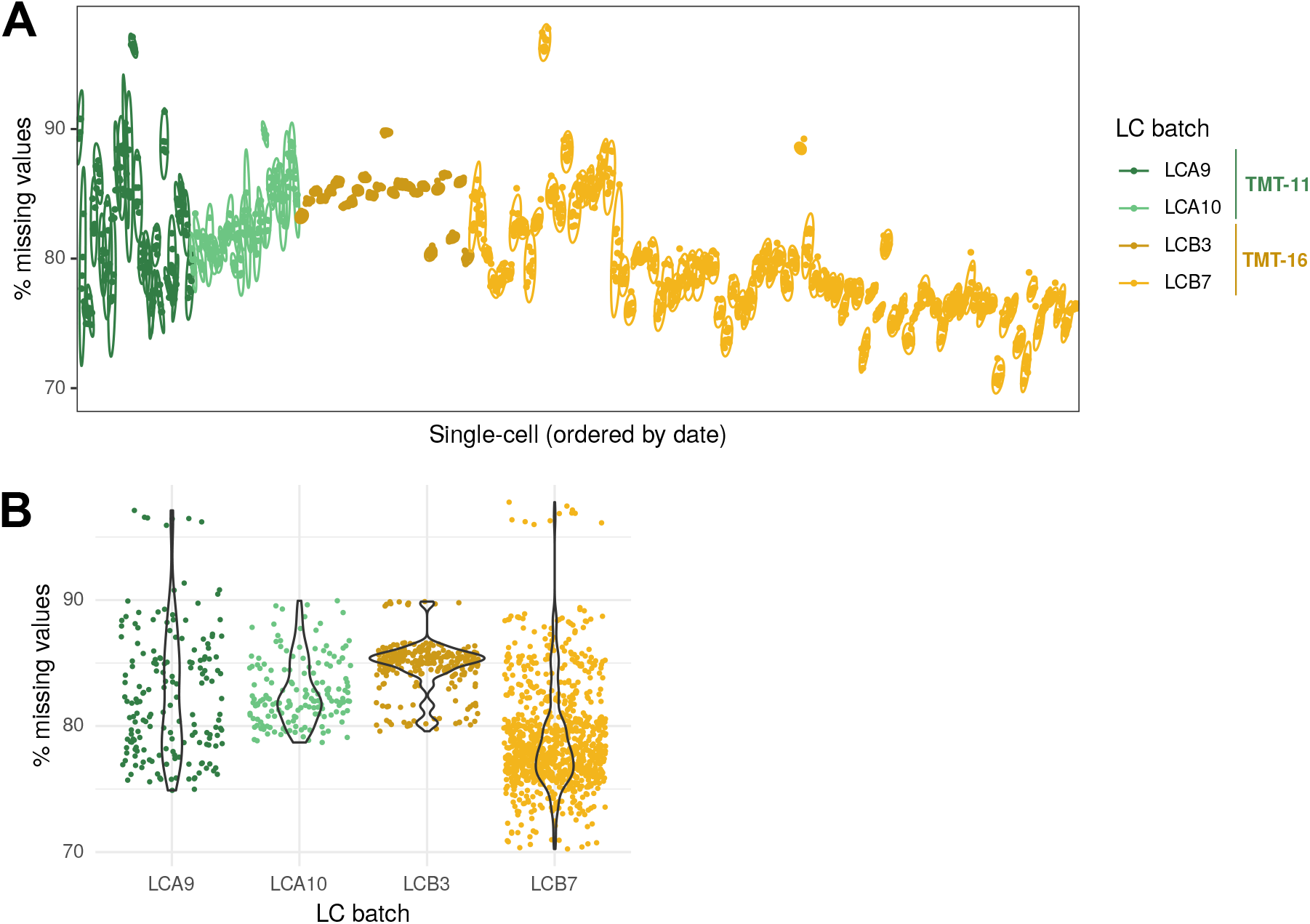
Influence of batch on data missingness. The proportion of missing data is shown for each single cell as a dot colored by LC batch. **A.** Effect of the MS run. Cells are ordered based on the acquisition date. The 95 % ellipses are drawn for every MS run. **B.** Effect of LC batch. Cells are grouped by LC batch. The missing data distribution within each batch is highlighted using violin plots.

A solution to this issue is to explicitly model the protein expression and the protein detection rate. The hurdle model is very compelling in this regard [37]. The hurdle model consist of two components. The first component is fit using the MSqRob model, that estimates peptide intensities as a function of sample covariates, and includes blocking factors for batch effect, taking into account the correlation between peptides belonging to the same protein [28]. Inference on the estimates allow to perform differential abundance analysis. The second component is a binary component that models the probability that an observation is missing as a function of sample covariates for each run independently. This second component is fit using a quasi-binomial regression and allows to assess differential detection. Another solution is provided by proDA [38], that constructs a sigmoidal probabilistic dropout model for each sample. This model is in turn used to infer means across samples and the associated uncertainty in the presence of missing values. Further research is needed to assess the performance of the model when applied to SCP data in the light of inflation of missing values, and to further adapt the algorithm to achieve principled SCP data analysis.

In conclusion, we believe there are two open paths of research that need to be explored to deal with the batch effect and data missingness challenge. First, we need to better understand the different mechanisms that influence missingness and batch effects in SCP data and how they differ from bulk proteomics. Benchmark data sets are therefore required to assess our ability to control for technical factors (e.g. operator, acquisition run, instrument, LC column, . . .) while preserving biological meaningful variability (e.g. cell type, cell state, treatment, . . .). Second, there is a need for dedicated models and methods that can disentangle the technical challenges that are batch effects and data missingness from the desired biological knowledge. `scp` represents an ideal environment for a standardized processing of the data and hence allowing comparison, integration and improvement of various existing methods available from other fields as well as benchmarking new methodological innovations.

## Acknowledgments

The authors would like to thank Nikolai Slavov, Edward Emmott and Harrison Specht for their openness in sharing all their data and scripts, and responsiveness in addressing questions and comments.

